# Crystal structure of *Anopheles* gambiae actin depolymerizing factor explains high affinity to monomeric actin

**DOI:** 10.1101/2024.06.16.598887

**Authors:** Devaki Lasiwa, Inari Kursula

## Abstract

Actin is an intrinsically dynamic protein, the function and state of which are modulated by actin-binding proteins. Actin depolymerizing factors (ADF)/cofilins are ubiquitous actin-binding proteins that accelerate actin turnover. Malaria is an infectious disease caused by parasites of the genus *Plasmodium*, which belong to the phylum Apicomplexa. The parasites require two hosts to complete their life cycle: the definitive host, or the vector, which is an *Anopheles* spp. mosquito, and a vertebrate intermediate host, such as humans. Here, the crystal structure of the malaria vector *Anopheles gambiae* ADF (*Ag*ADF) is reported. *Ag*ADF has a conserved ADF/cofilin fold with six central β-strands surrounded by five α-helices with a long β-hairpin loop protruding out of the structure. The G and F-actin binding sites of *Ag*ADF are conserved, and the structure shows features of potential importance for regulation by membrane binding and redox state. *Ag*ADF binds monomeric ATP- and ADP-actin with a high affinity, having a nanomolar K_d_, and binds to and effectively destabilizes actin filaments.

## Introduction

Actin is one of the evolutionarily most conserved proteins and the most abundant intracellular protein found in all eukaryotic cells. The dynamic remodeling of the actin cytoskeleton is involved in many biological functions, including motility, cell division, endocytosis, and intracellular trafficking (1). Actin cytoskeleton dynamics are regulated spatially and temporally through various actin-binding proteins (2). The actin depolymerizing factor (ADF)/cofilin family comprises severing proteins responsible for disassembly of filamentous actin (F-actin). All eukaryotes have ADF/cofilins, and they play critical roles in accelerating the actin cytoskeleton remodeling, thus affecting the dynamics of motile structures like lamellipodia (3), filopodia (4), *Listeria* comet tails (5), and neural growth cones (6). ADF/cofilins are also essential for the maintenance of contractile systems including contractile rings (7), stress fibers (8), and muscles (9) by modulating the quantity and length of actin filaments.

ADF/cofilins are small globular proteins that bind to the sides of the actin filaments usually with a preference for ADP-F-actin (3, 10). ADF/cofilin binding leads to severing of actin filaments, mainly at the pointed end of decorated and bare actin segments. ADF/cofilins can also bind to and accelerate depolymerization from the barbed ends of actin filaments when no ATP-bound globular actin (G-actin) is available. Full decoration of actin filaments by ADF/cofilin enhances depolymerization from the pointed and barbed ends (11). In addition, ADF/cofilins bind monomeric G-actin in a nucleotide dependent manner with a higher affinity to ADP-G-actin than to ATP-G-actin, and inhibit the rate of nucleotide exchange from ADP to ATP (12, 13). ADF/cofilins are regulated by multiple mechanisms to conduct their function in cells including phosphorylation/dephosphorylation (14, 15), variation in pH (16, 17), binding to phosphoinositides (18–20), and oxidation/reduction (21, 22).

Malaria is one of the most serious, life-threatening diseases, caused by unicellular eukaryotic apicomplexan parasites of the genus *Plasmodium.* The lifecycle of *Plasmodium* alternates between the definitive host or vector, which is a mosquito, and a vertebrate intermediate host, such as a human. Malaria is transmitted by the bite of infected mosquitos of the *Anopheles* genus. *Anopheles* spp. are abundant and widely distributed around the world. In tropical Africa, the most effective vector is *Anopheles gambiae* (23, 24)*. Plasmodium* spp. use an actomyosin-based mode of motility, termed gliding motility, to invade host cells (25, 26). Unlike other apicomplexan parasites, *Plasmodium* spp. express two isoforms of actin. Actin I is abundant and expressed in all life stages, whereas actin II is present only in the sexual stages within the mosquito (27, 28). In the malaria parasites, actin polymerization is strictly controlled by a limited set of regulators compared to other eukaryotic cells, and ADFs belong to the core set present in parasites. The two *Plasmodium* ADFs are substantially differ from each other and from higher eukaryotic counterparts (29).

Although apicomplexan host cell invasion is mainly powered by the parasite actomyosin glideosome complex (30), also modulation of the host actin cytoskeleton seems to be involved (31, 32). However, it is unclear whether the parasite and host actins and actin regulatory proteins come to contact *in vivo*. Curiously, we observed that *Plasmodium* actin II, which has functions in the mosquito stages of the parasite, copurifies with the insect *Spodoptera frugiperda* ADF/cofilin, and the complex is hard to disassemble. This serendipitous finding prompted us to characterize the malaria vector *Anopheles gambiae* ADF (*Ag*ADF). We describe here the first crystal structure of *Ag*ADF and its G- and F-actin related activities and compare its structure and biochemical functions with those of canonical and malaria parasite ADF/cofilins.

## Results

### *Ag*ADF has a canonical ADF fold with possible regulatory sites

Despite extensive research on ADF/cofilin family proteins, no structure has been available for *Ag*ADF. We determined the first crystal structure of *Ag*ADF at 1.68 Å resolution (Figure 1, Table 1, Figure S1). *Ag*ADF crystallized in space group P2_1_2_1_2 with two molecules in the asymmetric unit. The crystal had pseudo-translational symmetry, as indicated by the presence of a large off-origin peak in the Patterson map detected by Xtriage. Therefore, the R values are higher for this crystal structure than expected based on the resolution (33, 34) (Table 1). The electron density for most parts of the protein was clearly defined, so that the structure could be built with high confidence. Most side chain positions were unambiguous, except for some long side chains on the surface of the protein (Figure S1). The first three N-terminal and five C-terminal residues were not visible in the electron density and were therefore not built into the model.

**Figure 1:**
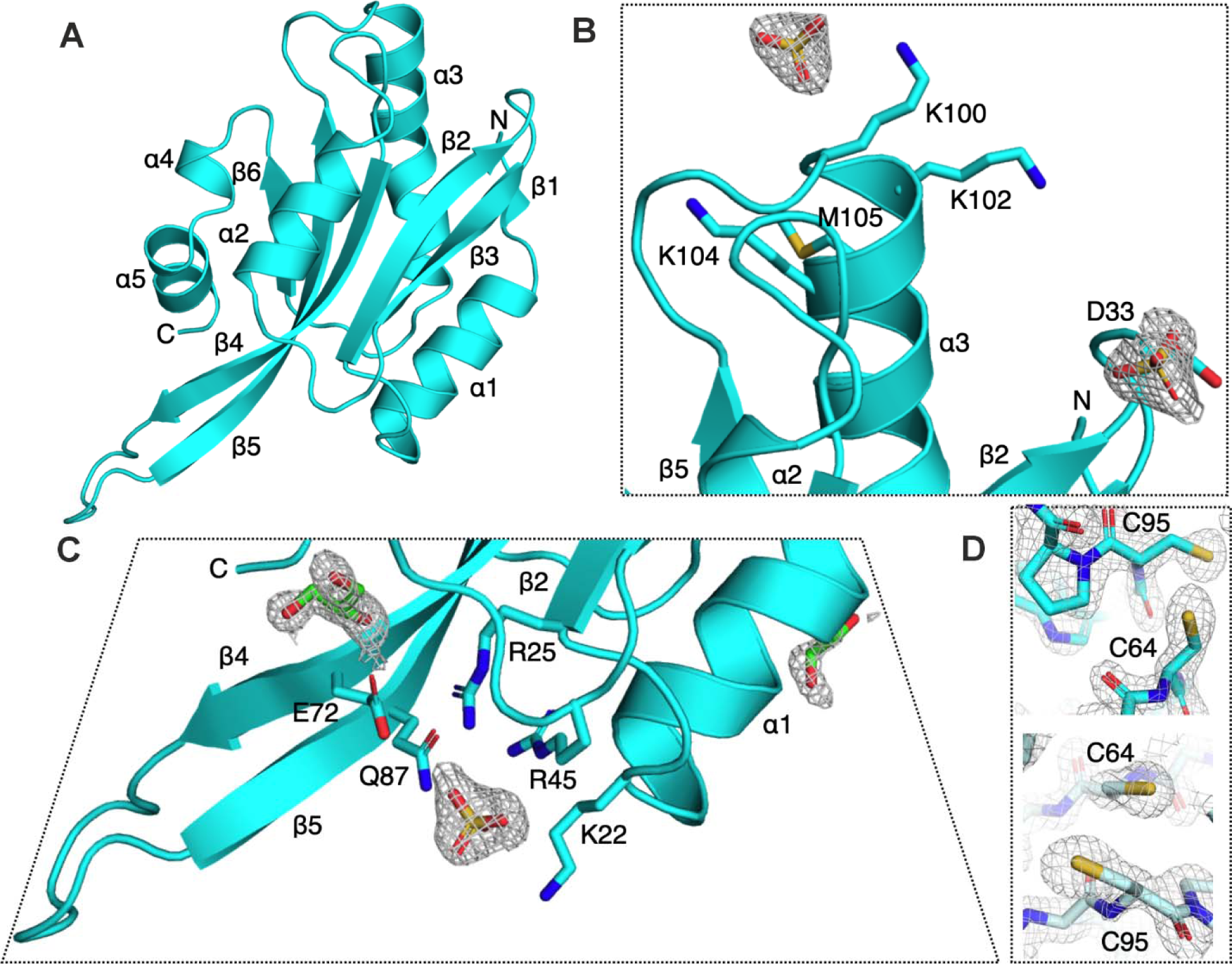
Crystal structure of *Ag*ADF. (A) A cartoon representation of the *Ag*ADF crystal structure with a central β-sheet sandwiched between five α-helices. The N- and C-termini, all α-helices and β-strands as well as the F-loop are labelled. (B) Sulphate ions bound close to the major G/F-actin binding site of *Ag*ADF. The sulphate ions and the side chains of residues near them are shown as sticks. The 2f_o_-f_c_ electron density, contoured at 1.5 σ around the sulphate ions is displayed. (C) The sulphate ions and glycerol molecules located near the F-loop. (D) Neither of the cysteine residues at close distance from each other, Cys-64 and Cys-95, in both molecules in the asymmetric unit form disulphide bonds in the crystal grown under reducing conditions. The 2f_o_-f_c_ electron density map is contoured at 1.5 σ.

**Table 1:** Crystallographic data collection and refinement statistics. Statistics for the highest resolution shell are shown in parentheses.

|  |  |
| --- | --- |
| Data collection | *AgADF* |
| Wavelength (Å) | 0.928 |
| Resolution range (Å) | 47.14–1.68 (1.74–1.68) |
| Space group | P2<sub>1</sub>2<sub>1</sub>2 |
| Unit cell dimensions |  |
| a, b, c (Å) | 62.42, 87.27, 56.01 |
| $\alpha$, $\beta$, $\gamma$ (°) | 90, 90, 90 |
| Total no. reflections | 444998 (23971) |
| Unique reflections | 35120 (3,066) |
| Multiplicity | 6.5 (4.8) |
| Completeness (%) | 98.47 (87.80) |
| $\langle I/\sigma(I) \rangle$ | 3.77 (0.84) |
| Wilson B-factor | 16.15 |
| R<sub>meas</sub> (%) | 35 (201) |
| CC<sub>1/2</sub> (%) | 99 (29.4) |
| **Model building and refinement** |  |
| No. of reflections | 35049 (3066) |
| R<sub>work</sub> | 0.2970 (0.4487) |
| R<sub>free</sub> | 0.3376 (0.4432) |
| No. of atoms |  |
| Protein | 2310 |
| Ligands* | 43 |
| Solvent | 104 |
| RMSD |  |
| Bonds (Å) | 0.014 |
| Angles (°) | 1.32 |
| Average B factors (Å<sup>2</sup>) |  |
| Protein | 29.01 |
| Ligands* | 42.09 |
| Water | 29.81 |
| Ramachandran plot |  |
| Favoured (%) | 98.55 |
| Allowed (%) | 1.45 |
| Outliers (%) | 0.00 |

\*Sulphate and glycerol molecules

*Ag*ADF has a typical ADF/cofilin fold (Figure 1A). The structure consists of a central mixed beta sheet (β2-β6) surrounded by α-helices (α1-α5). The central part of the β-sheet formed by strands β3-β2-β4-β5 is antiparallel, while the short β1 at the N-terminus and β6 at the opposite edge of the sheet are parallel to their neighboring strands. The five main α-helices flank either side of the central β-sheet, with α1 and α3 on one side and α2 and the broken C-terminal helix (α4-α5) on the opposite side. The structure shows conserved features of the G/F-site and F-site, which include the characteristic long helix α3 (G/F site), the β4-β5 loop called the F-loop (F-site), and the C-terminal helix (F-site) (Figure 1).

ADF/cofilins are regulated by membrane phosphoinositide binding, which inhibits their interaction with actin (16, 31). In the *Ag*ADF crystal structure, there are altogether three sulphate binding sites per monomer that may mimic the phosphoinositide-binding sites (Figure 1B and C). One sulphate is bound to the G/F-site at the N-terminal end of α3, coordinated by Lys-100 (Figure 1B). This sulphate is located on a symmetry axis and is, thus, bound to the same site in both monomers in the asymmetric unit. Another sulphate is also shared between two monomers in the crystal and is bound in one monomer at the stem of the F-loop, coordinated by Lys-22, Arg-25, Arg-45, Glu-72, and Gln-87, and in another monomer close to the G/F-site, more loosely coordinated by Asp-33 (Figure 1B and C).

ADF/cofilins are susceptible to oxidation/reduction of cysteine residues, which is a regulatory factor. Under oxidizing conditions, cofilin alters its cellular location, particularly its accumulation in mitochondria (21, 22, 35). *Ag*ADF has four cysteines (Cys-11, Cys-64, Cys-77, and Cys-95). Of these, only Cys-77 is solvent-exposed and lies in the F-loop. Cys-64 occupies a position similar to Cys-80 in *Homo sapiens* cofilin *(Hs*Cof) (36), and its side chain is at close distance to that of Cys-95. However, there is no disulphide bond to be observed in either monomer in the crystal (Figure 1D), which is not surprising given that the reducing agent tris (2-carboxyethyl) phosphine (TCEP) was used throughout purification and crystallization. Cys-11 is buried and has no potential disulphide pair. Thus, there are two possible ways that oxidation could contribute to the regulation of *Ag*ADF under oxidizing conditions: modification or intermolecular disulphide formation *via* Cys-77 or intramolecular disulphide formation between Cys-64 and Cys-95).

### *Ag*ADF has several conserved residues of the ADF/cofilin family including a long F-loop

*Ag*ADF shares 36% and 37% sequence identity with *Saccharomyces cerevisiae* cofilin (*Sc*Cof), and *Arabidopsis thaliana* ADF1 (*At*ADF1) respectively, for which crystal structures are known. There are several highly conserved residues in the protein family (Figure 2). The conserved region at the N-terminus includes residues Ser-3, Gly-5, and two hydrophobic residues.

**Figure 2:**
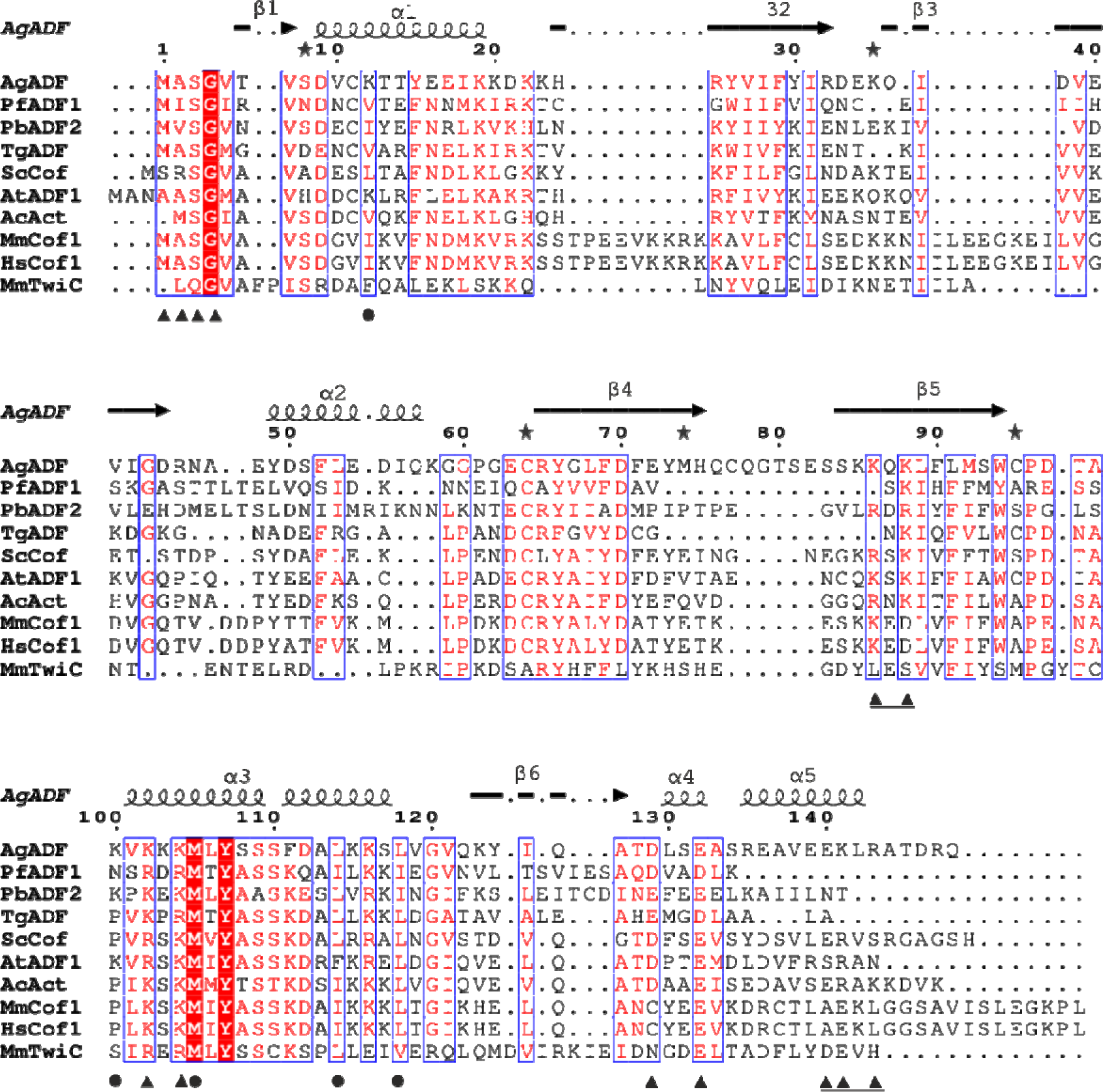
Multiple sequence alignment of *Ag*ADF and selected other ADF/cofilins. The amino acid sequence of *Ag*ADF was aligned with other ADF/cofilin family members using ClustalW2 (60). Strictly conserved residues are shown in red boxes and regions of residues with similar properties are indicated with blue boxes. The secondary structure elements of *Ag*ADF are shown and named above the alignment. G-actin-binding sites identified in yeast cofilin by mutagenesis (19) and synchrotron footprinting (81) are marked with black triangles and circles, respectively. Residues involved in the F-actin-binding site are marked with underlined black triangles. The sequences include those of *P. falciparum* ADF1 (*Pf*ADF1), *P. berghei* ADF2 (*Pb*ADF2), *A. gambiae* ADF (*Ag*ADF), *T. gondii* ADF (*Tg*ADF), *S. cerevisiae* cofilin (*Sc*Cof), *A. thaliana* ADF1 (*At*ADF1), *A. castellanii* actophorin (*Ac*Act), *M. musculus* cofilin-1 (MmCof), *H. sapiens* cofilin (*Hs*Cof), and *M. musculus* Twf-C (*Mm*TwiC).

ADF/cofilins are regulated by phosphorylation of residue Ser-3, which inhibits their interaction with actin (37, 38). Ser-3 is not resolved in the crystal structure because the N-terminal residues are disordered, and electron density can be observed only from Gly-4. Other conserved regions are within the β4 strand, including Asp-70 and four hydrophobic residues, and from Lys-102 to Asp-112, of which Met-105 and Tyr-107 are present in all ADF/cofilin family members, including *Ag*ADF. Highly conserved residues located in the hydrophobic core of ADF/cofilins such as Tyr-66, Trp-94, Pro-96 and Tyr-107 are important for protein stability or folding (39).

The root means square deviations (RMSD) for the Cα positions between *Ag*ADF and *Sc*Cof (PDB ID: 1COF) (40), and between *Ag*ADF and *At*ADF1 (PDB ID: 1F7S) are 1.1 Å and 1.4 Å, respectively (Figure 3A). The Cα RMSD values for the *Ag*ADF when superimposed on the malaria parasite *Plasmodium falciparum* ADF1 (*Pf*ADF1) is 1. 9 Å and on *Plasmodium berghei* ADF2 (*Pb*ADF2) 1.1 Å (Figure 3B). The F-loop in *Ag*ADF is longer than in other ADF/cofilins. The connecting loop between α2 and β4 is longer in *Ag*ADF. Compared to the malaria parasite ADFs, the differences to *Pf*ADF1 are much larger*, Pf*ADF1 lacks a protruding F-loop and the C-terminal helix α5 (Figure 3B). The differences to *Pb*ADF2, which resembles the canonical ADFs, are smaller and comparable to the other ADF/cofilins used for comparison. However, α2 of both *Plasmodium* ADFs and α4 of *Pb*ADF2 are longer than those in *Ag*ADF and other ADF/cofilins (Figures 2 and 3B).

**Figure 3:**
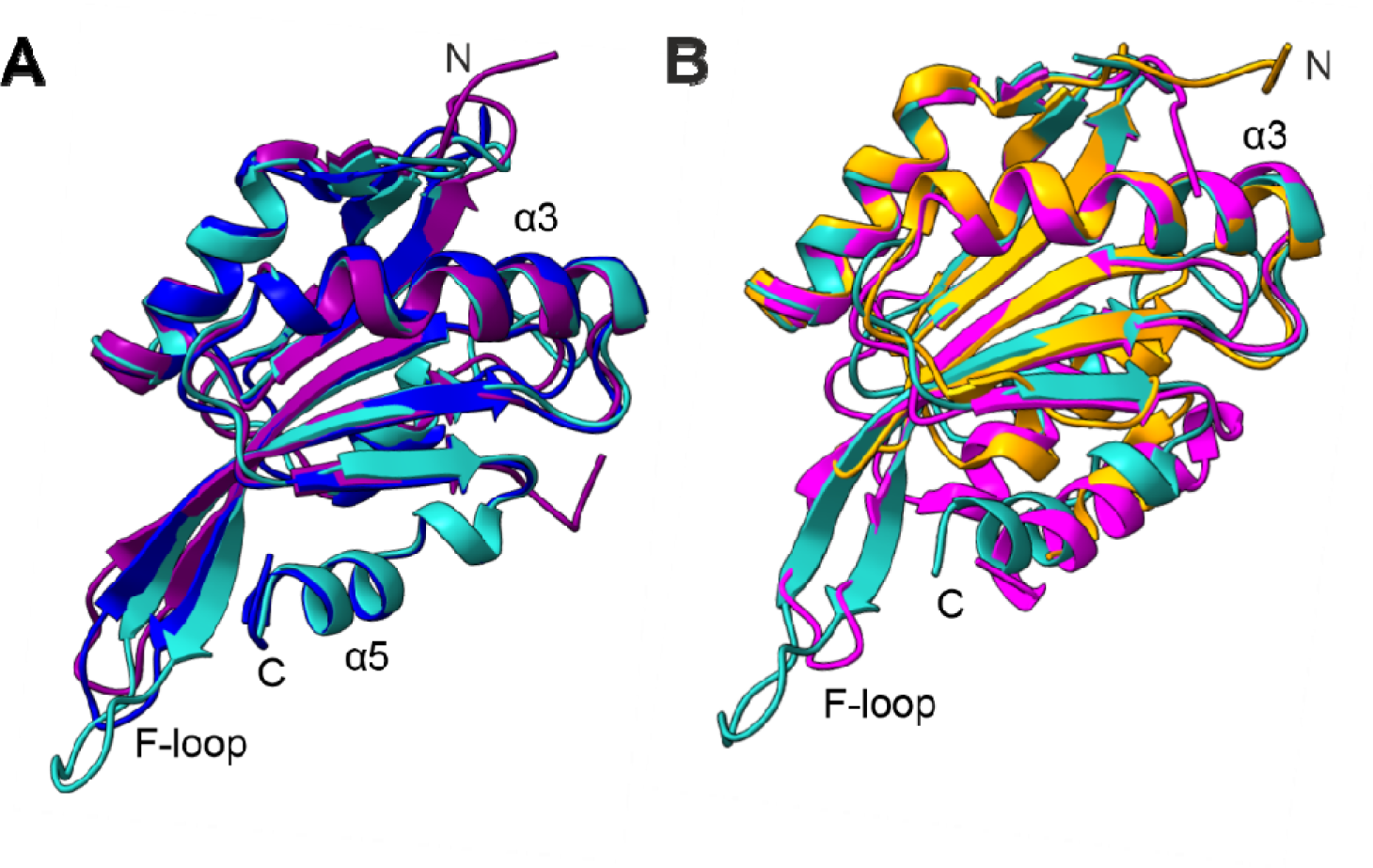
Structural comparison of *Ag*ADF with other members of the ADF/cofilin family. (A) Superposition of *Ag*ADF (cyan) with *Sc*Cof [PDB ID: 1COF (40); blue], and [*At*ADF1 PDB ID: 1F7S (82); purple]. The N- and C-termini, α3, α5, and the F-loop are labelled. (B) Superposition of *Ag*ADF (cyan) with *Pf*ADF1 [PDB ID: 2XF1 (29); orange] and *Pb*ADF2 [PDB ID: 2XFA(29); magenta]. The N- and C-termini, α3 as well as the F-loop are labelled.

### *Ag*ADF binds G-actin with high affinity

ADF/cofilins have two regions for actin binding. These are the G/F-site and the F-site (41). The G/F-site is responsible for binding both G-actin and F-actin, whereas the F-site is required for binding to F-actin and for F-actin severing activity. To get insight into the binding of the *Ag*ADF to G-actin, we generated a model of *Ag*ADF in complex with *Gallus gallus* actin using AlphaFold (42) (Figure 4A). The predicted structure is overall similar to the crystal structure of mouse twinfilin C (Twf-C) with rabbit muscle α-actin (43). Three major sites are involved in the G-actin interaction: the N-terminus, the long α3, and the C-terminal helix. The N-terminal sequence formed by Ser, Gly, and hydrophobic residues of ADF/cofilin is conserved in *Ag*ADF. This region is flexible in the *Ag*ADF crystal structure and lacks secondary structure. In the AlphaFold model of the *Ag*ADF-actin complex, Met-1, Val-3, and Gly-4 are involved in the interaction with actin (Figure 4A). The long α3 forms a major actin-binding site and inserts into a groove between subdomains (SDs) 1 and 3 of G-actin. Two highly conserved basic residues (Arg-267 and Arg-269 in Twf-C) in α3, are directly involved in actin interactions. The corresponding residues in *Ag*ADF are lysines (Figures 2). In the model, these residues do not interact with each other, but Lys-102 and Ser-350 (actin) are close to each other. The hydrophobic residues around the basic residues Val-101, Met-105, and Leu-106 interact with actin, similarly to the interaction in the Twf-C actin complex. Asp-129, Glu-132, and Glu-140 in the C-terminal helix are involved in interaction with actin in addition to Gln-126, which has similar interaction as Glu-296 in the Twf-C actin complex (Figure 4A).

**Figure 4:**
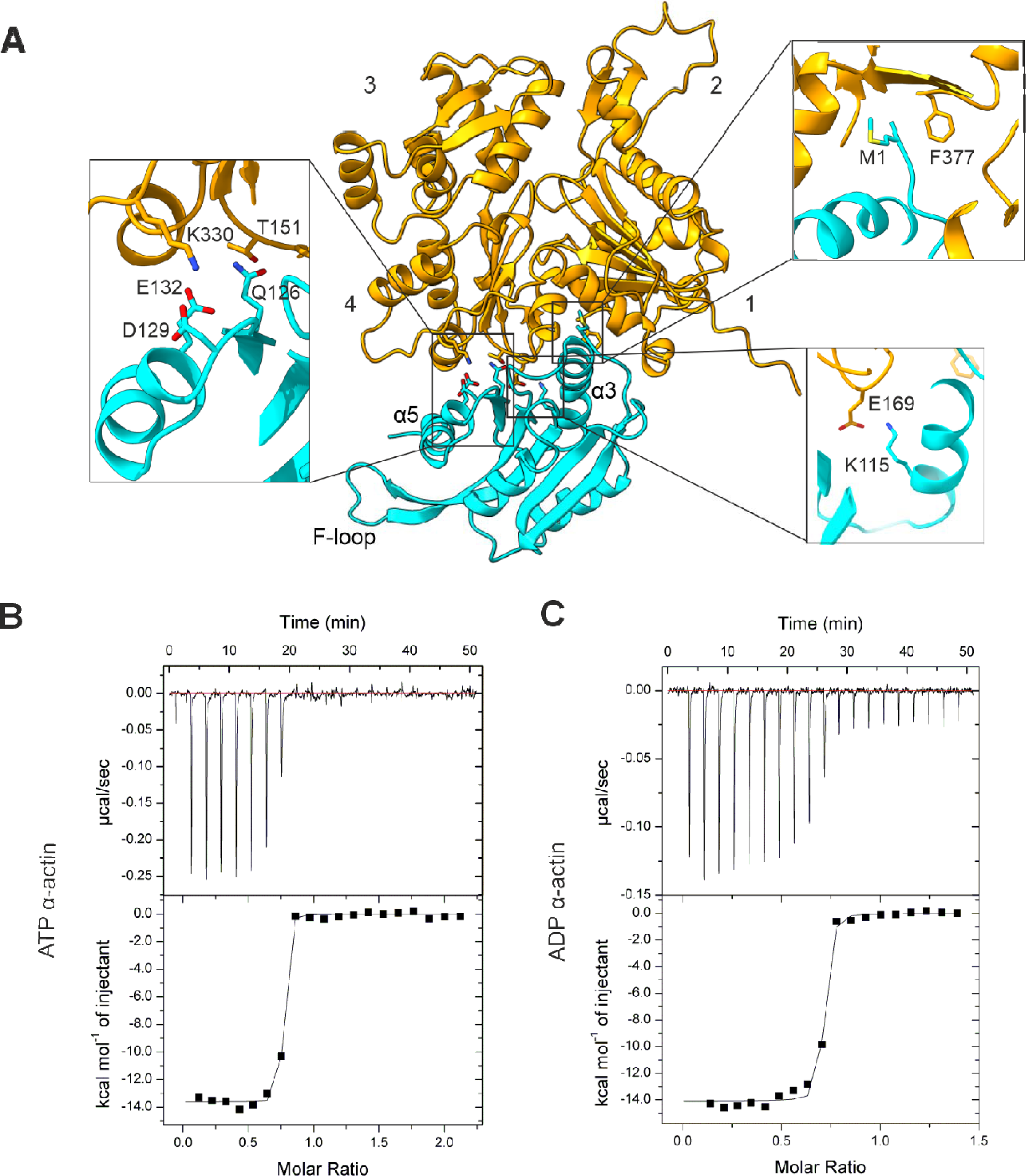
*Ag*ADF interaction with G-actin. (A) Model of the *Ag*ADF (cyan) and *G. gallus* G-actin (yellow) complex generated using AlphaFold (40). The zoomed-in views show the interactions discussed in the text. The actin SDs 1-4 are labelled as are the α3, α5, and F-loop of the *Ag*ADF. (B) ITC of *Ag*ADF G-actin in the presence of ATP. (C) ITC of *Ag*ADF G-actin in the presence of ADP. In both (B) and (C), the negative peaks indicate an exothermic reaction. The area under each peak represents the heat released after the injection of *Ag*ADF into the G-actin solution (upper panel). Binding isotherms were obtained by plotting the peak areas against the ADF/G-actin molar ratio. The lines represent the best-fit curves obtained from the least square regression analysis, assuming a one-site binding.

The structural features of *Ag*ADF described above support binding to G-actin. To test this, isothermal titration calorimetry (ITC) was performed under low ionic conditions, where actin stays in monomeric form. The titration of ADP-G-actin and ATP-G-actin with *Ag*ADF reveals a stoichiometry close to 1 in both cases (Figure 4B and C). The dissociation constant (K_d_) values were 0.8 and 1.6 nM for ATP-G-actin and ADP-G-actin, respectively, with ΔH of −13.6 ± 0.1 kcal mol^-1^ and -TΔS 1.2 kcal mol^-1^ for ATP-G-actin and ΔH of −14.1 ± 0.1 kcal mol^-1^ and -TΔS 2.1 kcal mol^-1^ for ADP-G-actin. These observed high affinity with a nanomolar K_d_ between *Ag*ADF and G-actin in the presence of either ADP or ATP, thus, correlates with the structural features of the binding sites in the crystal structure.

### *Ag*ADF binding to F-actin

The effect of *Ag*ADF on the kinetics of actin assembly was measured using actin labeled with fluorescent N-(1-pyrene) iodoacetamide (here referred to as pyrene). The fluorescence of pyrene F-actin is approximately 25-fold higher than that of monomeric pyrene G-actin (44). Here, 0.5, 1, 2, and 4 μM *Ag*ADF were incubated with α-actin, of which 5% of actin was labelled with pyrene, and the fluorescence intensity after initiating polymerization was measured over time (Figure 5A). *Ag*ADF increased the initial nucleation (Figure 5B), but then inhibited elongation (Figure 5C) and final steady state levels (5D) of actin polymerization at all concentrations tested. At a 1:1 *Ag*ADF-to-actin ratio, polymerization was almost completely inhibited. Cosedimentation assays were used to characterize the ability of *Ag*ADF to bind and disassemble actin filaments. Both *Ag*ADF and G-actin alone remained in the supernatant fraction after centrifugation at 100000 g (Figure 5E). *Ag*ADF cosedimented with F-actin at all the concentrations tested and significantly reduced the amount of actin in the pellet as compared to the F-actin control without *Ag*ADF (Figure 5E and F).

**Figure 5:**
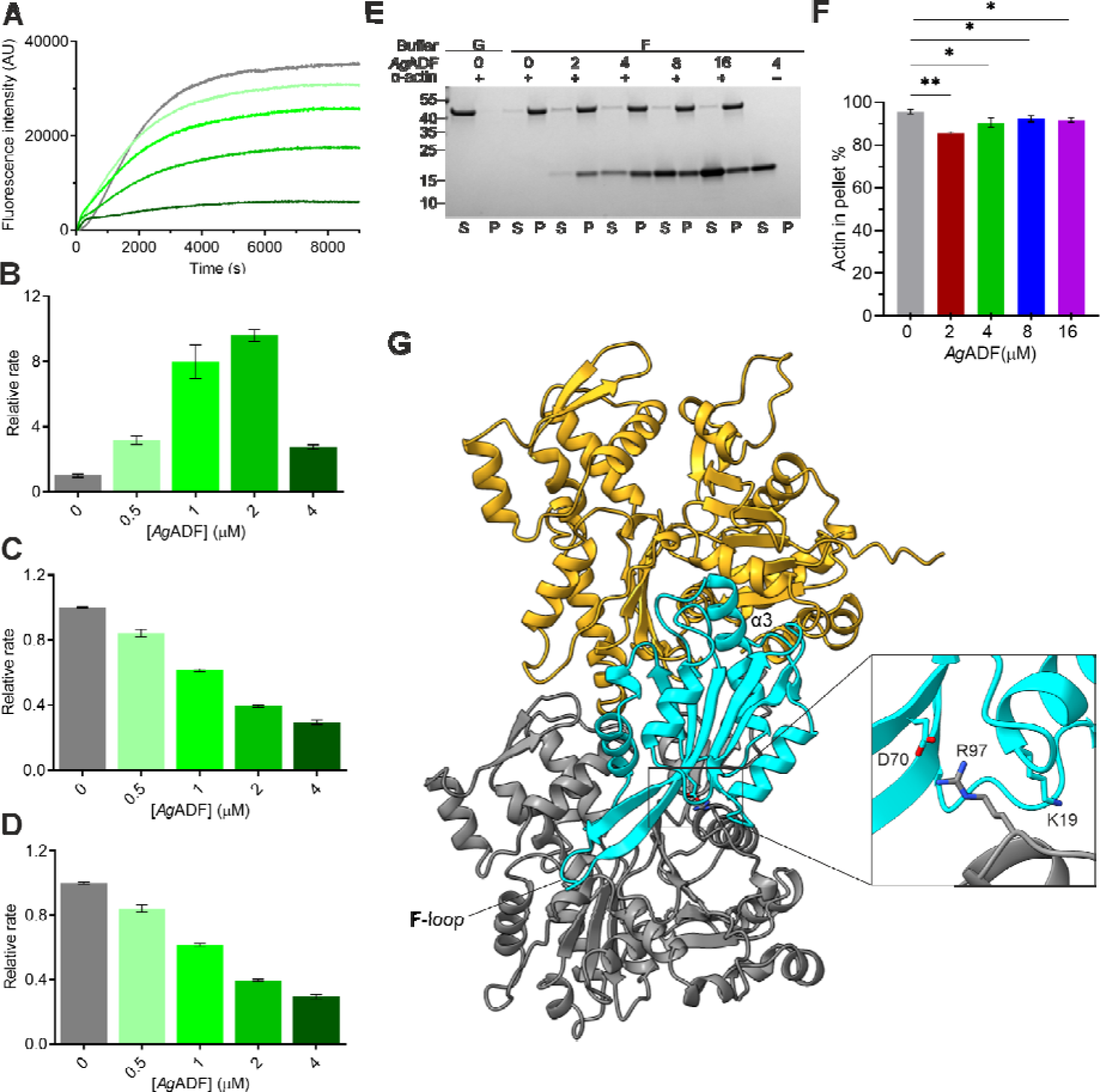
*Ag*ADF interaction with F-actin. (A) Polymerization curves of 4 μM pyrene-labeled α-actin alone and in the presence of 0.5-4 μM *Ag*ADF. (B) Relative initial rates determined from the first 65 s, representing the nucleation phase. (C) Relative initial rates determined from 500 s, representing the linear elongation phase. (D) Relative steady state values determined from the time frame of 7000-8000 s. The data are presented as mean ± standard deviation and n = 3. (E) Sedimentation of α-actin in the absence and presence of 2-16 μM *Ag*ADF. A representative gel is shown. The actin concentration in each sample was 4 µM, and the *Ag*ADF concentrations are displayed in µM above the gel images. G and F represent G-buffer and F-buffer, respectively. S and P denote the supernatant and pellet, respectively. The molecular weights of relevant standards in kDa are shown on right side of the gel. (F) Quantification of the proportion of actin in pellet fractions in the co-sedimentation assay. The error bars represent mean ± standard deviation and n = 3. Asterisks represent statistical significances determined using unpaired two-tailed t-tests for the actin in pellet, where *P < 0.05 and **P < 0.01 (G) Model of the complex between *Ag*ADF and *G. gallus* a longitudinal F-actin dimer generated using Alphafold (40). The two actin protomers are shown as gray and yellow and *Ag*ADF as cyan. The zoomed-in view shows a salt bridge between Asp-70 (*Ag*ADF) and Arg-97 (actin) at the interface. AgADF α3 and F-loop are labelled.

The F-actin binding site in ADF/cofilins extends from the G/F site through the C-terminus to the F-loop on the opposite side of the protein and interacts with two actin protomers in the filament. These binding interfaces observed in the high-resolution chicken cofilin-actin cryo-EM structure are to a large extent conserved in *Ag*ADF, but the F-loop is longer than in other family members (45). To gain insight into actin filament binding, we generated an AlphaFold model of *Ag*ADF-bound to a longitudinal actin dimer as in the filament (Figure 5G). In addition to the shared G/F-actin binding interface on one protomer, *Ag*ADF shows an F-actin binding site, which interacts with SDs 1 and 2 of the longitudinally adjacent actin subunit, similarly to mammalian cofilins. The F-loop and C-terminus of *Ag*ADF are longer and comprise most of the F-actin binding sites. In addition to these, Lys-19, Asp-70, and Asp-97 also interact with actin (Figure 5G). These residues are close to Lys-22, Glu-72, and Lys-100, which interact with two of three bound sulphates.

### *Ag*ADF conformation in solution

In parallel with the crystal structure determination, small-angle X-ray scattering (SAXS) was used to determine the size, shape, and oligomeric state of *Ag*ADF in solution. The purified recombinant *Ag*ADF was folded and globular in solution, as indicated by the scattering curve (Figure 6A) and the dimensionless Kratky plot (Figure 6B). The maximum interatomic distance of approximately 60 Å (Figure 6C) as well as the radius of gyration (Rg) of approximately 18 Å, the Porod volume, and the calculated molecular weight (Table 2) are consistent with a monomer in solution. The *Ag*ADF crystal structure fits well into the SAXS envelope, confirming its monomeric state and showing that the crystal structure represents the overall structure and conformation in solution (Figure 6D). Compared to *Pf*ADF1, *Ag*ADF has a more elongated structure in solution, similar to *Pb*ADF2 (46).

**Figure 6:**
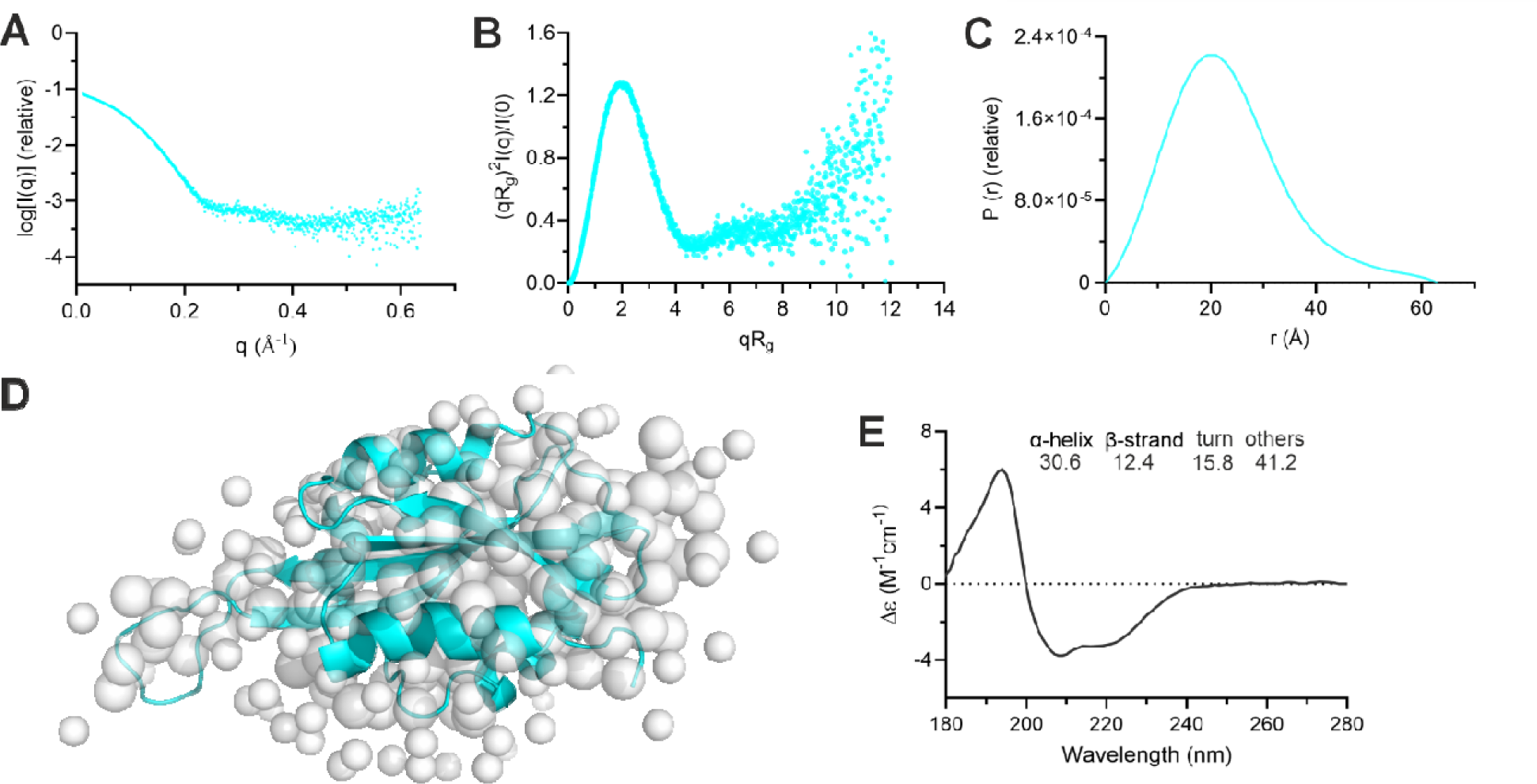
Solution structure of *Ag*ADF. (A) X-ray scattering curve of *Ag*ADF. (B) Dimensionless Kratky plot. (C) Real space distance distribution function. (D) An *ab initio* model generated using GASBOR overlaid with the crystal structure of *Ag*ADF. (E) SR-CD spectra of *Ag*ADF. The distributions of the major secondary structure components are shown in the inset as percentages.

**Table 2:** The estimated SAXS invariants and modelling parameters of *Ag*ADF.

| Sample | AgADF |
| --- | --- |
| I(0) (absolute, Guinier) | 0.072 |
| I(0) (absolute, real space) | 0.070 |
| R<sub>g</sub> (Å, Guinier) | 18.98 |
| R<sub>g</sub> (Å, real space) | 18.01 |
| D<sub>max</sub> (Å) | 62.91 |
| V<sub>porod</sub> (nm<sup>3</sup>) | 276 |
| MW (kDa), theoretical | 17.06 |
| MW (kDa), SAXS | 16.12 |
| Gasbor $\chi^2$ | 2.17 |

Synchrotron radiation circular dichroism (SR-CD) spectroscopy was used to determine the secondary structure contents of *Ag*ADF in solution (Figure 6E). Deconvolution of the SR-CD spectra indicated 31% α-helix, 12% β-strand, 16% turn, and 41% other structure, as calculated using data from 180 to 250 nm using the BeSTSel server (47). Except for the β-strand contents, this agrees with the secondary structure contents calculated from the *Ag*ADF crystal structure (Table S1).

## Discussion

The ADF/cofilin family proteins are multifaceted cellular players (48). They regulate actin filament dynamics through G- and F-actin binding, depolymerization, F-actin severing, G-actin sequestering activity, and by controlling the rate of nucleotide exchange in actin monomers (49–52). Many structures have been determined of ADF/cofilins from mammals, yeast, and Apicomplexa. All these share the highly conserved ADF-homology fold that is also observed in destrin, gelsolin, and twinfilin (29, 36, 40). *A. gambiae* is one of the most efficient vectors of the malaria parasite. Here, the *A. gambiae* ADF/cofilin homologue, *Ag*ADF, was characterized structurally and functionally.

*Ag*ADF has the conserved ADF/cofilin fold with an N-terminal flexible region, a long α3 helix, and a long F-loop, which projects out from the structure. Sequence comparisons and modeling of *Ag*ADF in a complex with G-actin (Figures 2 and 4A) demonstrated that the G-actin binding site of *Ag*ADF is conserved, including the N-terminus, a long α3, the turn connecting strand β6, and the C-terminal helices (43). The N-terminus of the *Ag*ADF is structurally mobile and disordered in the crystal structure. It is highly conserved in the ADF/cofilin proteins, particularly Ser-3, which is an important contact site for interactions with actin and a phosphorylation target (15, 38). The positively charged residues in α3, which interact with G-actin, are conserved in *Ag*ADF. Similar low nanomolar (2-30 nM) K_d_s to G-actin as seen for *Ag*ADF (Figure 4B and C) have also been reported for mouse ADF/cofilins (cofilin-1, cofilin-2, and ADF) and ADP-G-actin (53). The K_d_ values for the interaction of the *Toxoplasma gondii* ADF (*Tg*ADF) and the ADF/cofilins from *Trypanosoma brucei* cofilin with ADP-G-actin, also determined by ITC, were 20 nM and 80 nM, respectively (54, 55). ADF/cofilin proteins typically bind ADP-actin 10-100-fold higher affinity than the ATP-actin (53, 56). Contrary to this, we did not observe a large difference in the affinity of *AgA*DF to ADP-G-actin or ATP-G-actin, and the affinity was even slightly higher to ATP-G-actin. The high affinity of *Ag*ADF for ATP-G-actin may explain the why *Ag*ADF inhibits overall polymerization *via* sequestration of monomers in non-polymerizable complexes with *Ag*ADF. However, *Ag*ADF increases initial nucleation rate, which is probably due to its severing activity, making more free ends available before the monomer sequestering comes into play.

In conventional ADF/cofilins, the F-site includes a pair of basic residues in the F-loop and charged residues at the C-terminus of the protein (49, 57). These residues are conserved in *Ag*ADF. For example, Lys-86 and Lys-88 in the F-loop, corresponding to Arg-80 and Lys-82 of yeast cofilin; Glu-140 and Lys-141 in the C-terminal helix of *Ag*ADF corresponding to Glu-134 and Arg-135 of yeast cofilin (Figure 2). These F-loop basic residues and the C-terminal charged residues increase the stability of the interaction of ADF/cofilins with F-actin (37, 57). Consistent with this, 2 μM *AgADF* cosedimented nearly quantitatively with F-actin (polymerized from 4 µM G-actin). In *Ag*ADF, the F-loop that projects out of the structure is longer than that of other ADF/cofilins, being for example five residues longer than in *Sc*Cof. However, the stabilizing interaction with F-actin was associate with a limited net disassembly of actin filaments. These two properties, stable interaction and disassembly, are inversely related. Similar activity has been seen with *Caenorhabditis elegans* UNC-60B (57). ADF/cofilins showing stable interaction with F-actin may be more effective at filament severing (57).

Actin binding of ADF/cofilins is typically inhibited by phosphoinositides (19, 20, 58). A recent study suggested that ADF/cofilins interact with phosphoinositide headgroups through a large positively charged protein surface. The positively charged surface is created by a cluster of highly conserved residues, including Lys-95, Lys-96, Lys-112, Lys-114, Lys-125, Lys-126, Lys-127, Lys-132, and His-133 in the case of *Hs*Cof (18). Sequence alignment of *Ag*ADF with *Hs*Cof showed that these lysine residues are conserved in *Ag*ADF (Figure 2). In addition, sulphate molecules present in the crystal structure interact with Lys-100 and Glu-87, which closely correspond to Lys-112 and Lys-114 of *Hs*Cof. These residues overlap with the G-actin binding site. In support of this, the PPM Webserver (59) for positioning proteins in membranes predicted similar binding interactions with the membrane (Figure S2).

In summary, *Ag*ADF adopts a conserved ADF/cofilin fold with conserved binding motifs for both G- and F-actin binding. Consistent with this, it binds G-actin with a nanomolar K_d_ and cosediments with and depolymerizes F-actin. Surprisingly, *Ag*ADF binds both ATP and ADP actins with very similar affinities. Considering the high affinity of *Ag*ADF to α-actin, future structure determination of actin-*Ag*ADF complexes could be attempted. Actin II, which is specific for the sexual stages of the malaria parasite within the mosquito host, copurifies with insect ADF *in vitro*. Thus, further *in vivo* and *in vitro* studies on whether actin II binds to *Ag*ADF might be an interesting future line of research.

## Experimental procedures

### Sequence alignment of *Ag*ADF with other ADF/cofilin proteins

A multiple sequence alignment of AgADF and other ADF/cofilin proteins was generated with ClustalW2 (60) and visualized using ESPript (61). UniProtKB accession numbers were as follows: *Ag*ADF, *A. gambiae* (A0NGL9); *At*ADF1, *A. thaliana* ADF1 (Q39250); *Ac*Act, *Acanthamoeba castellanii* actophorin (P37167); *Sc*Cof, *S. cerevisiae* cofilin (Q03048); *Mm*Cof, *Mus musculus* cofilin-1 (P18760); HsCof, *Homo sapiens* cofilin (P23528); *Pf*ADF1, *P. falciparum* ADF1 (Q8I467); *Pb*ADF2, *P. berghei* ADF2 (Q3YPH0); and *Tg*ADF, *T. gondii* ADF (B9Q2C8).

### Protein expression and purification

*Ag*ADF with an N-terminal His_6_ tag cloned into a pETNKI-his-SUMO3 vector (NKI Protein Facility, Amsterdam, Netherlands) was constructed by Mr. Markku Soronen. For expression, the construct was transformed into *E. coli* Rosetta (DE3) (Novagen, Darmstadt, Germany). The SUMO3 domain located between the tag and the ADF ensured a native N-terminus, as the sentrin-specific protease 2 used for cleavage of the tag, leaves no extra amino acids in the N-terminus. Selected transformants were inoculated into Luria Bertani medium at 37 with 50 µg/mL kanamycin and 34 µg/mL chloramphenicol and grown overnight at 37°C. Expression cultures were grown in ZYM-5052 autoinduction medium (62) at 37 for 4 h after inoculation with 1% preculture. The cultures were then cooled to 20 and left for 60 h. The cells were harvested by centrifugation at 5020 g for 45 min, washed with phosphate-buffered saline and either stored at −20 or used directly for purification.

A two-step purification protocol, involving immobilized metal affinity chromatography followed by size exclusion chromatography (SEC) was used to purify *Ag*ADF from fresh or frozen cell pellets. The cell pellets were resuspended in a lysis buffer containing 20 mM Tris-HCl (pH 8.0), 50 mM NaCl, 5 mM imidazole, 5 mM 2-mercaptoethanol (β-ME), supplied with one tablet of ethylenediaminetetraacetic acid free Sigma*FAST* protease inhibitor cocktail tablet (Merck KGaA, Darmstadt, Germany) per 100 ml buffer. The cells were lysed by sonication using a Branson 450 Digital Sonifier (Marshall Scientific LLC, Hampton, NH, USA) with a 1/2” tapped disruptor horn for 2 min with 30 s pulses. The lysate was clarified by centrifugation at 42500 g for 30 min. The supernatant was applied onto equilibrated HisPur nickel-nitrilotriacetic acid (Ni-NTA) matrix (Thermo Fisher Scientific Inc., Waltham, MA, USA) and incubated for 1 h under gentle agitation. The resin was washed extensively with wash buffer [20 mM Tris-HCl (pH 8.0), 50 mM NaCl, 20 mM imidazole, 5 mM β-ME] and eluted with elution buffer [20 mM Tris-HCl (pH 8.0), 50 mM NaCl, 300 mM imidazole, β-ME]. The eluted protein was treated with sentrin-specific protease 2 and dialyzed against 20 mM Tris-HCl (pH 8.0), 50 mM NaCl, 5 mM β-ME. The sample was then re-incubated with freshly equilibrated Ni-NTA matrix for 1 h, and the flow-through was collected. The matrix was washed with the dialysis buffer complemented with 15 mM imidazole to prevent unspecific binding to the matrix and to maximize the yield. The flow-through and washes were then pooled and concentrated using a concentrator with a 3 kDa molecular weight cut-off (Millipore, Burlington, MA, USA) and the protein was filtered through polyvinylidene fluoride ultra-free membrane filter with a 0.22 μm pore size (Millipore). The concentrated and filtered sample was then finally passed through a HiLoad 10/300 Superdex 75 column (GE Healthcare, Chicago, IL, USA) equilibrated with SEC buffer [20 mM Tris-HCl (pH 8.0), 50 mM NaCl, 0.5 mM TCEP]. The peak fractions were pooled, concentrated, filtered, snap-frozen, and stored at −70 until further use.

Acetone powder from pig (*Sus scrofa*) sirloin muscle was prepared and α-actin purified using an established protocol (63). The protein was stored either as G-actin under dialysis in G-buffer (10 mM HEPES pH 7.5, 0.2 mM CaCl2, 0.5 mM ATP and 0.5 mM TCEP) or F-actin on ice.

### Isothermal titration calorimetry

ITC was performed at 25 with a stirring rate of 750 rpm using a MicroCal iTC 200 calorimeter (GE Healthcare). Stock solutions of α-actin and *Ag*ADF were dialyzed extensively against 5 mM Tris-HCl (pH 7.5), 0.2 mM CaCl_2_, 0.2 mM ATP or 0.2 mM ADP, 0.5 mM TCEP. 8 µM α-actin in the cell and 80 µM *Ag*ADF in the syringe after centrifugation and degassing were used in the binding titrations. An aliquot of 280 µL of α-actin was loaded into the cell and titrated with 40 µL *Ag*ADF. Titrations for binding were initiated by one injection of 0.4 µL followed by 3.6 µL injections with 150 s between injections to allow baseline recovery. Eleven injections were monitored. Each titration was performed in duplicate. *Ag*ADF was also titrated to buffer as a control under similar conditions to account for the heat of dilution. The data were analyzed using ORIGIN software (OriginLab Corporation, Northampton, MA, USA). The first injection was excluded from the analysis. A curve fit using a one-set-of-sites fitting model was used to determine the K_d_, stoichiometry of binding, and enthalpy change (ΔH) for all interactions.

### Circular dichroism spectroscopy

Secondary structural compositions of *Ag*ADF were determined using SR-CD. SR-CD spectra for *Ag*ADF at 0.12 mg/mL were recorded three times in water between 170-280 nm in a Hellma cylindrical absorption cuvette (Suprasil quartz, Hellma GmbH & Co. KG, Müllheim, Germany) with a pathlength of 0.5-1 mm at the AU-CD beamline at the ASTRID2 synchrotron (ISA, Aarhus, Denmark) at 25. Buffer spectra were subtracted, and CD units converted to Δε (M^-1^cm^-1^) using CDtoolX (64). Secondary structure deconvolutions were carried out using BeStSel (65).

### Small-angle X-ray scattering

SAXS was used to determine a low-resolution solution structure of *Ag*ADF in order to get insight on its size, shape, and oligomeric state in solution. Purified *Ag*ADF was dialyzed overnight in SEC buffer at 4. SAXS data were collected from samples at 5-15 mg/mL at the CoSAXS beamline at MAX IV ring (Lund, Sweden). The data were further analyzed using PRIMUS (66) and programs of the ATSAS package (67). The resulting models were visualized using PyMOL 2.0 (Schrödinger, NY, USA). *Ab initio* models generated by GASBOR (68) are shown.

### Crystallization, data collection and refinement

Initial crystallization screening for *Ag*ADF was carried out using following commercial screens: JCSG-plus, PACT premier, Proplex (all from Molecular Dimensions, Holland, OH, USA), Crystal screen I and II (Hampton Research, CA, USA), and home-formulated salt grid and factorial screens (69). Crystals grew at 4 in a condition with 0.17 M ammonium sulphate, 25.5% w/v polyethylene glycol 4000, 15% v/v glycerol in the JCSG-plus screen at 20 mg/mL concentration. Crystals picked from this drop were directly frozen in liquid nitrogen before shipping to the synchrotron facility for data collection.

Four different data sets from *Ag*ADF crystals were collected on the I04-1 beamline at Diamond Light Source (Oxfordshire, UK). The data were processed and scaled using XDS (70). A CCP4-MTZ file format file was generated using XDSCONV, and molecular replacement was performed using PHASER (71) within the PHENIX suite (72). Only one data set gave a solution. The model was manually built using Coot (73, 74), and refinement of the structure using data up to 1.68 Å was carried out using PHENIX.refine (75). The analysis of the model was performed using Coot (73) and the ChimeraX 1.4 (76). Structural comparisons between *Ag*ADF and other ADF/cofilin members were performed using PDBeFold (77).

### Actin polymerization assays

For polymerization assays, actin labelled with pyrene, as described previously (78) was used. The polymerization assays were performed according to an established protocol (78). In brief, polymerization of 5% pyrene labelled 4 µM α-actin preincubated with varying concentrations (0.5 - 4 µM *Ag*ADF) in G-buffer was initiated by adding 10×F-buffer to final concentrations of 50 mM KCl, 4 mM MgCl_2_, 1 mM EGTA to a total reaction volume of 150 µl. The increase in fluorescence upon polymerization was followed for 2 h using an Infinite M1000 Pro (Tecan) multimode plate reader at 25 with excitation and emission wavelengths of 365 nm (9 nm bandwidth) and 407 nm (20 nm bandwidth), respectively. The reactions were carried out in triplicate. Data were exported and analyzed using Graphpad Prism 8 (GraphPad Software, La Jolla, CA, USA). All polymerization curves were set to start from zero fluorescence intensity. Relative rate and Plateau levels of the polymerization curves were determined as average values from the range of 65-200 s, 500-1000 s and 7000-8000 s respectively. The experiment was repeated three times using different protein batches.

### Actin co-sedimentation assays

An actin co-sedimentation assay was performed to assess the interaction of ADF with α-actin. A total of 4 µM α-actin in G-buffer was polymerized for 2 h at room temperature in the presence of 2-16 µM *Ag*ADF in a total volume of 100 µL. Polymerization was achieved by adding 10×F-buffer to final concentrations of 50 mM KCl, 4 mM MgCl_2_, and 1 mM EGTA. The samples were then subjected to high-speed ultracentrifugation (100000 g for 1 h) at 20 using an Optima TL-100 benchtop ultracentrifuge (Beckman Coulter, Indianapolis, IN, USA). *Ag*ADFs and α-actin alone were processed identically as a controls. The supernatants and pellets were separated, and the pellets was resuspended in 100 µL G-buffer. Both fractions were mixed with 25 µL of 5 x sodium dodecyl-sulphate polyacrylamide gel electrophoresis (SDS-PAGE) sample buffer [250 mM Tris-HCl (pH 6.8), 10% SDS, 50% glycerol, 0.02% Bromophenol Blue and 1.43 M β-ME]. The samples were incubated for 5 min at 95°C, and 10 µL of each sample was analyzed on 4-20% SDS-PAGE gels. Protein bands were visualized using PageBlue staining (Thermo Fisher Scientific Inc.). The gels were imaged using a ChemiDoc XRS+ system (Bio-Rad, Hercules, CA, USA). Gel bands were quantified using IMAGEJ (79). The assay was repeated three times using different protein batches.

### Protein membrane interaction study

The position of the *Ag*ADF on the membrane was estimated using the positioning of Proteins in Membrane (PPM) server version 3.0 (59). *Ag*ADF coordinate was submitted to the PPM server. Calculations used mammalian plasma membrane excluded heteroatoms, water, and detergents. The *Ag*ADF is declared by PPM program as a peripheral protein.

## Data availability

All the plasmids and relevant data used to support the findings of this study are available upon request from the corresponding authors. The coordinates and structure factors have been submitted to the Protein Data Bank under the accession code (9FP8).

## Supporting information

This manuscript contains supporting information (80).

## Acknowledgments

We thank Dr. Matti Myllykoski for collecting the X-ray diffraction data and Dr. Maiju Uusitalo and Oda Caspara Krokengen for collecting the SAXS data. We acknowledge the use of the I04-1 beamline on the Diamond Light Source (Oxfordshire, UK) for diffraction data collection, the AU-CD beamline on ASTRID2 at ISA (Aarhus, Denmark) for SR-CD measurements, and the CoSAXS beamline at MAX IV (Lund, Sweden) for access to SAXS beamtime. Access to the facilities and the expertise of the Biocenter Oulu Proteomics and Protein Analysis as well as Structural Biology core facilities, members of Biocenter Finland, are gratefully acknowledged. This work was funded by the Sigrid Jusélius Foundation (I.K.), the Academy of Finland (I.K), the Emil Aaltonen Foundation (I.K), the Jane and Aatos Erkko Foundation (I.K.), and the Norwegian Research Council (I.K.).

## Author contributions

D.L. and I.K. conceptualization; D.L. methodology; D.L. and I.K. formal analysis and validation; D.L. writing original draft; I.K. writing – review and editing. I.K. supervision, project administration, and funding acquisition.

## Abbreviations

ADF: actin depolymerizing factor
Ag: Anopheles gambiae
Ac: Acanthamoeba castellanii
Act: actophorin
At: Arabdidopsis thaliana
β-ME: beta mercaptoethanol
Cof: cofilin
EDTA: ethylenediaminetetraacetic acid
Hs: Homo sapiens
K_d_: dissociation constant
ITC: isothermal titration calorimetry
Mm: Mus musculus
NiNTA: nickel nitrilotriacetic acid
PBS: phosphate-buffered saline
Pb: Plasmodium berghei
Pf: Plasmodium falciparum
Rg: radius of gyration
RMSD: root mean square deviation
SAXS: small-angle X-ray scattering
Sc: Saccharomyces cerevisiae
SDS-PAGE: sodium dodecyl-sulphate polyacrylamide gel electrophoresis
SEC: size exclusion chromatography
SR-CD: synchroton radiation circular dichroism
TCEP: tris(2-carboxyethyl)phosphine
Tg: Toxoplasma gondii
Twf-C: twinfilin C

