## Supplemental materials for "Crystal structure of *Anopheles* gambiae actin depolymerizing factor explains high affinity to monomeric actin"

Materials included:

Figure S1

Figure S2

Table S1

**Figure S1**

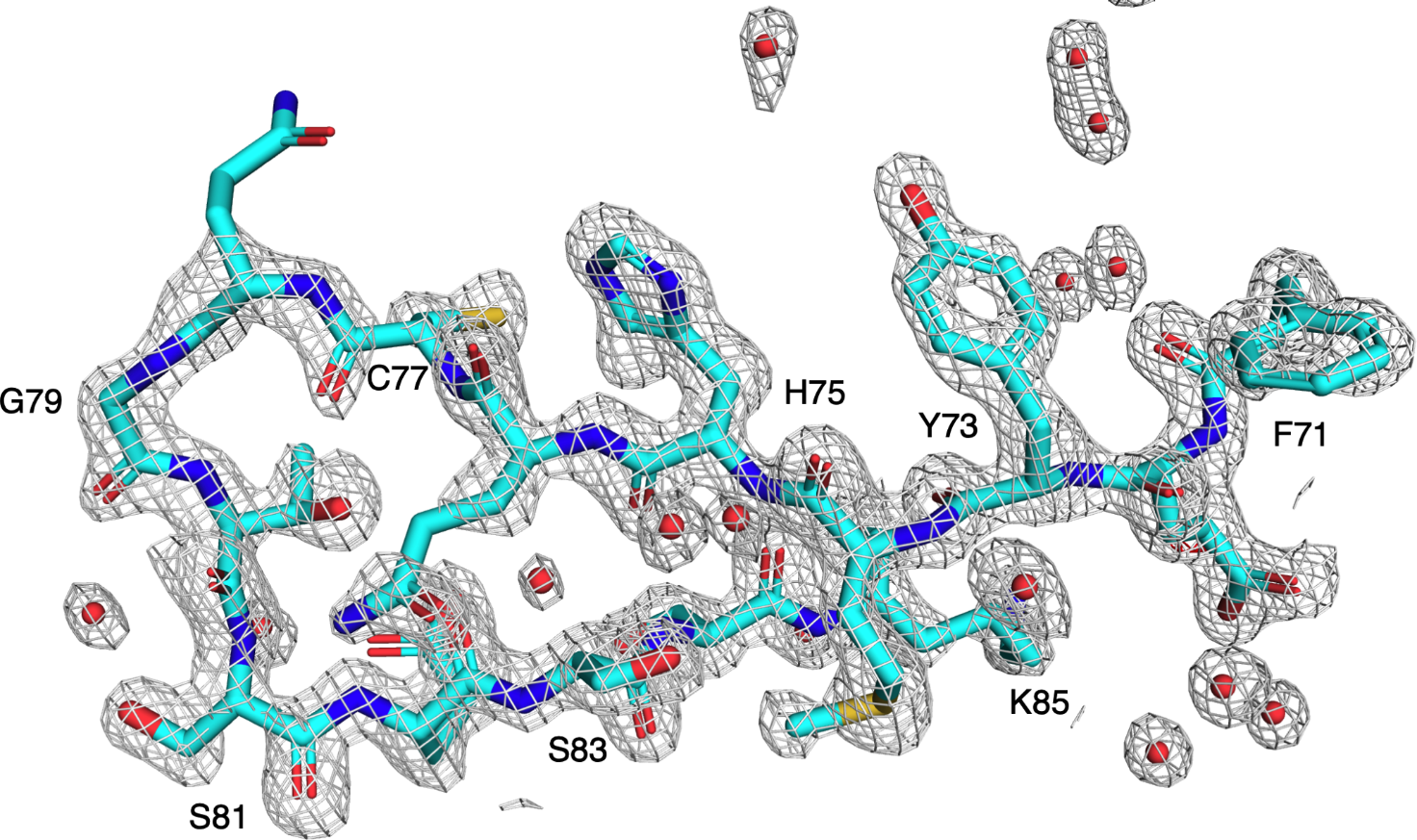

**Figure S1:** Representative electron density around the F-loop (residues 71-85, labeled). The 2f_o_-f_c_ map (gray) is contoured at 1.5 σ. The f_o_-f_c_ map is contoured at +/- 5 σ (blue and red, respectively, no peaks visible). Water molecules are shown as red spheres.

**Figure S2**

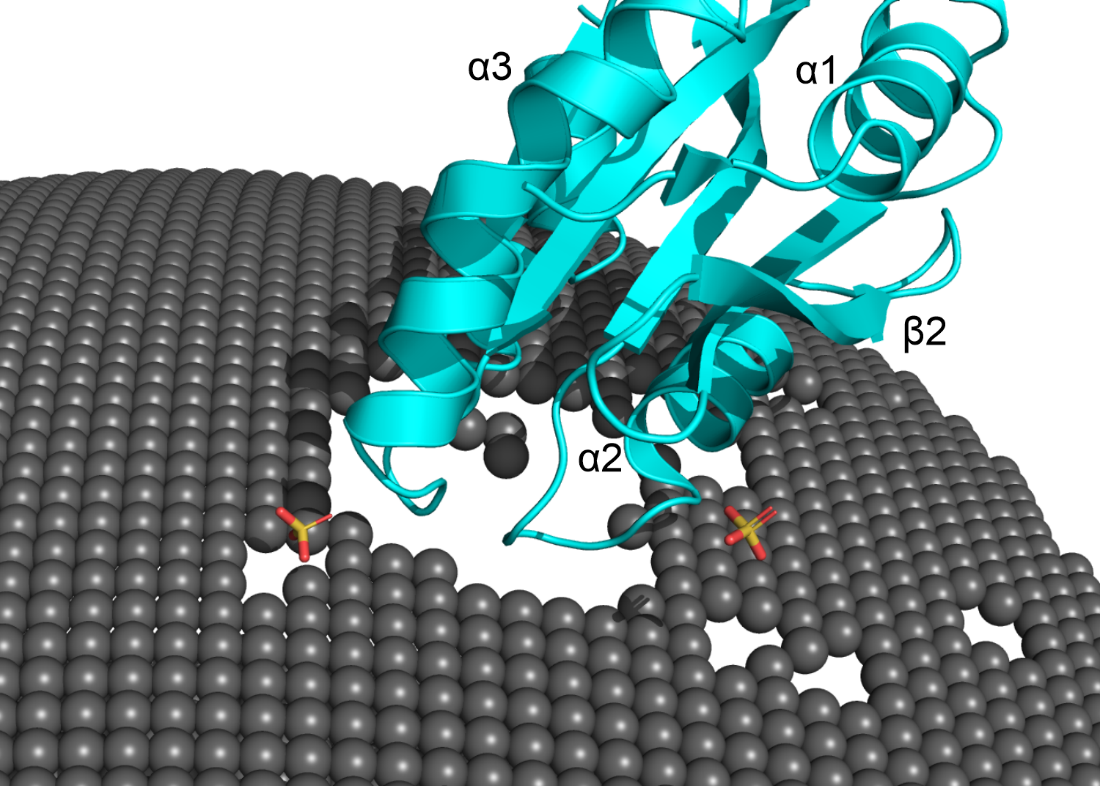

**Figure S2:** Model of *Ag*ADF (cyan) interacting with a membrane. The bound sulphate ions are shown as sticks. The secondary structure elements α1, α2, α3, and β2 of *Ag*ADF are labelled.

**Supplementary table S1**

Secondary structure contents (%) calculated from the *Ag*ADF crystal structure using PDBMD2CD (72).

| α-helix | 32.4 |
| --- | --- |
| β-sheet | 33.1 |
| turn | 14.2 |
| others | 20.2 |
